# Using recent baselines as benchmarks for megafauna restoration places an unfair burden on the Global South

**DOI:** 10.1101/2021.03.13.435233

**Authors:** Sophie Monsarrat, Jens-Christian Svenning

## Abstract

The potential for megafauna restoration is unevenly distributed across the world, along with the socio-political capacity of countries to support these restoration initiatives. We show that choosing a recent baseline to identify species’ indigenous range puts a higher burden for megafauna restoration on countries in the Global South, which also have less capacity to support these restoration initiatives. We introduce the Megafauna Index, which considers large mammal’s potential species richness and range area at country-level, to explore how the responsibility for megafauna restoration distributes across the world according to four scenarios using various temporal benchmarks to define species’ indigenous range – current, historical (1500AD), mid-Holocene and Pleistocene. We test how the distribution of restoration burden across the world correlates to indicators of conservation funding, human development, and governance. Using a recent or historical baseline as a benchmark for restoration puts a higher pressure on African and southeast Asian countries while lifting the responsibility from the Global North, where extinctions happened a long time ago. When using a mid-Holocene or Pleistocene baseline, new opportunities arise for megafauna restoration in Europe and North America respectively, where countries have a higher financial and societal capacity to support megafauna restoration. These results contribute to the debate around benchmarks in rewilding initiatives and the ethical implications of using recent baselines to guide restoration efforts. We suggest that countries from the Global North should reflect on their responsibility in supporting global restoration efforts, by increasing their support for capacity building in the South and taking responsibility for restoring lost biodiversity at home.

## Introduction

Overcoming the current biodiversity crisis will require ambitious conservation solutions to not only protect nature, but also re-expand nature, through increased ecosystem restoration (Leclère et al. 2020) —a vision in line with the recently declared 2021-2030 UN Decade on Ecosystem Restoration. The Intergovernmental Science-Policy Platform on Biodiversity and Ecosystem Services (IPBES) defines restoration as *“any intentional activity that initiates or accelerates the recovery of an ecosystem from a degraded state”* (IPBES 2018). This definition asks the question of which reference baseline should be considered as the predegraded state and used as an appropriate benchmark for restoration. Is the appropriate turning point in human-induced changes the 19th century Industrial Revolution, the 16th century start of European expansion, or even the Late Pleistocene dispersal of *Homo sapiens* throughout the world? What are the socio-political implications of choosing a common baseline to guide restoration efforts in different regions of the world? Answering these questions requires an understanding of the long-term interaction between natural processes and human-induced changes and how these interfere with the financial and sociopolitical capacity of countries to support restoration.

Biodiversity is not uniformly distributed in space, which results in countries having complementary roles in attempting to safeguard biodiversity and limit species extinctions (5Strassburg et al. 2020). Different countries also have different capacities and resources when it comes to nature management and conservation. Many of the highest biodiversity conservation priority areas are in some of the world’s poorest countries (Brooks et al. 2006), where the pressure for conservation is high but financial resources are low (Miller et al. 2013, Reed et al. 2020). This creates tension between the need to meet global conservation targets and the gap in conservation expenditure in some areas (Balmford et al. 2003). To meet conservation objectives and reduce the rate of biodiversity loss, increased support from local, national and global communities is needed, often requiring a transfer of resources from north to south (Balmford and Whitten 2003), commensurate with the number of threatened species in recipient countries (Miller et al. 2013). These aid flows remain the largest source for conservation in the low- and middle-income countries of the developing world and are positively associated with a reduction in the rate of biodiversity loss (Waldron et al. 2017). Whether these resources are appropriately distributed according to the pressure for future ecosystem restoration remains to be investigated.

An increasing number of studies support a strong connection between rewilding as a restoration strategy and sustainable human development, noting that the promotion of rewilding in policy and decision-making would strongly support the post-2020 biodiversity goals (Perino et al. 2019, Pereira et al. 2020, Svenning 2020, Pasgaard et al. Under revision). Trophic rewilding (Svenning et al. 2016) recognizes the key role that megafauna species have on ecosystems processes, from seed dispersal, frugivory, herbivory and nutrient cycles (Pérez-Méndez et al. 2016, Villar et al. 2020, Enquist et al. 2020)— to effects on fauna and flora communities (Galetti et al. 2017), vegetation structure (Sandom et al. 2014, Riesch et al. 2020) and carbon storage (Bello et al. 2015). The long-history of coevolution with extant species and the disproportionate impact they have on the environment (Enquist et al. 2020) make megafauna species a potentially key element in our efforts to restore biodiverse and resilient ecosystems (Svenning et al. 2016, Fernández et al. 2017, Perino et al. 2019, Schowanek et al. 2021). While megafauna species can have significant cultural (Feldhamer et al. 2002, Berti et al. 2020), social (Maller et al. 2019) and economic (Lindsey et al. 2007) value to humans, their conservation may also come at a high financial (Lindsey et al. 2018) and societal (Jordan et al. 2020) cost. Countries’ contribution to the conservation of the world’s terrestrial megafauna is unevenly distributed today (Lindsey et al. 2017), with 40 of the most severely underfunded countries for conservation containing 32% of all threatened mammalian diversity (Waldron et al. 2013). Identifying the potential relative contribution of countries to global rewilding efforts in the future strongly depends on how we identify the set of megafauna species that could be reintroduced in each country.

Variations in rewilding practices relate to the choice of temporal benchmark to guide restoration, which requires deciding on a baseline geography to understand the potential range of species, where populations could recolonize naturally or be restored in the future. Most global biodiversity initiatives turn to pre-European landscapes (often using the arbitrary date of 1492 when Columbus arrived in the New World) or pre-industrial conditions of the mid-19^th^ century as a benchmark for the delineation of species’ indigenous range and the reference against which to compare current ecosystem condition (Keulartz 2016, Akçakaya et al. 2018, Stephenson et al. 2019). However, these baselines are biased with a Eurocentric perspective that overlooks variations in the timing of human impacts across the world (Sanderson 2019), which can lead to arbitrary and potentially unfair restoration. For megafauna species, extinctions in the Late Pleistocene and early Holocene 50 000 to 7 000 years ago – mostly concentrated in the Americas and Australia, and to a lesser extent in Eurasia (Stuart 2015, Smith et al. 2018) – as well as historical (Monsarrat et al. 2019, Teng et al. 2020) and recent declines (Ripple et al. 2015, 2016) have heavily reshaped global distribution patterns (Faurby and Svenning 2015). Hence, considering current, relictual, patterns of distribution risks hindering our understanding of optimal habitat for species (Kerley et al. 2020), underestimating areas where it could be restored (Monsarrat et al. 2019), and ultimately leading to less ambitious restoration objectives (Balmford 1999).

Paleoecological approaches to rewilding science seeks to push back this historical horizon, to better comprehend the ecological dynamics of a prehuman world and inform future conservation interventions (Lorimer et al. 2015). Pleistocene systems have been suggested as a relevant “pre-human” ecological benchmark to inform rewilding, as they describe how ecosystems functioned independently of modern humans. This recognizes that the pre-extinction Pleistocene fauna, which was the norm 50 000 years ago – a blink of an eye in evolutionary times – represents a functional state that had been typical for millions of years (Stegner and Holmes 2013, Smith et al. 2018). Pleistocene rewilding, though controversial (Rubenstein et al. 2006) is promoted by conservation biologists in North America (Donlan et al. 2006), South America (Galetti 2004) and Russia (Zimov 2005). In Europe, the “pre-agrarian” ecological conditions of the early to mid-Holocene ~6-10 000 years ago have often been used as a benchmark to guide rewilding efforts in Europe, with more emphasis on the introduction of large herbivores, to reinstate functional ecosystem dynamics via the restoration of resource-driven bottom-up systems (Keulartz 2016). Choosing one of these baselines as a benchmark for rewilding will affect which megafauna species are classified as native from an area, and thus worth considering for restoration.

Here, we aim to understand how the choice of fixed temporal baselines as benchmarks to guide restoration can lead to arbitrary and potentially unfair restoration targets for megafauna restoration in certain parts of the world. How does the “burden” for megafauna restoration distributes across the world according to the benchmark used to define species’ indigenous range? And how does that correlate to socio-economic contexts and the capacity of countries to reach these restoration objectives? We hypothesize that choosing a common, recent, baseline as a benchmark for global megafauna restoration puts a higher burden in locations with higher poverty, low institutional capacity and weak social safety nets. We map the distribution of restoration burden under four different scenarios using different temporal baseline to define species’ indigenous range and examine how this correlates to countries’ governance, development levels and conservation funding.

## Material and methods

### Species ranges and baseline scenarios

We define megafauna species as terrestrial mammals at least 45kg in body mass for herbivores and 21.5kg for carnivores, following Malhi et al. (2016), excluding *Homo* species. For each species, we mapped current, historical (1500AD) and present-natural ranges. Present-natural ranges can be viewed as pre-*Homo sapiens* ranges adjusted for present-day climate (Faurby and Svenning 2015). They were originally derived by Faurby & Svenning (2015) using current IUCN extent of occurrence for species unaffected by human pressure and by combining current IUCN ranges with historical distributions of species, fossil co-occurrence and known range modifications caused by humans. Current IUCN extent of occurrence and present-natural range maps were extracted from the PHYLACINE database (v1.2.1, Faurby et al. 2018, 2020). Detailed information about these range maps can be found in the metadata associated with this database (Faurby et al. 2018, 2020). For three species *(Panthera pardus, Helarctos malayanus, Ursus arctos),* we mapped historical ranges as provided by the IUCN Red List by using the “Extinct” range, defined as the area where the species was formerly known or thought very likely to occur post 1500 AD (IUCN Red List Technical Working Group 2019). For other species, we reviewed the literature for information on pre-historical extinctions within biogeographical realms and clipped areas where the species is known to have gone extinct before 1500AD from the present-natural range. This coarse spatial resolution prevents detailed mapping of local extinctions patterns but is sufficient for the coarse resolution at which this analysis is performed. We rasterized all range maps to a resolution of 96.5 km × 96.5 km at ± 30° latitude using the Behrmann equal-area projection.

Using these range maps, we simulated the following four restoration scenarios: a) *Current,* where all extant megafauna species are present in their current range; b) *Historical,* where all extant megafauna species are restored within their historical range (baseline: 1500AD); c) *Holocene,* where all extant megafauna species are restored within their present-natural range, d) *Pleistocene,* where all megafauna species (both extant and recently extinct species) are restored in their present-natural range. All globally extinct species were assigned to the Pleistocene baseline, even if a subset survived into the early Holocene (e.g. Crees and Turvey 2014, van der Plicht et al. 2015), so the Holocene scenario corresponds to a mid-Holocene temporal baseline. The Pleistocene scenario is not realistic since it includes globally extinct species, but it is as a counterfactual to explore the question: how would the burden of large mammal restoration be distributed if no extinctions had occurred since the Late Pleistocene? It also allows for the possibility of restoring some of the evolutionary and ecological potential that was lost in the Late Pleistocene, by using surrogate species as functional analogues to replace extinct species (Donlan et al. 2006) or managing already established non-native functional analogues of extinct megafauna (Lundgren et al. 2020).

### Species richness and Megafauna Index

Species richness was calculated by counting the number of species extant in each country under each restoration scenario. A species was considered extant in a country if any part of its range polygon overlapped with the country’s polygon. Species richness alone does not appropriately reflect the level of effort required to restore megafauna communities. For example, a species has the same weight whether it occurs in very small portion or throughout the entire country. We present the Megafauna Index (MI), which better considers the distribution of species within each country. MI combines the number of megafauna species in each country and the cumulative proportional area that they occupy within it. It is calculated using the following formula: 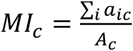, where *a_ic_* is the area of the range for species’ *i* within country *c* in *A_c_* km^2^ and *Ac* is the area of country *c* in km^2^. As an example, a value of MI=10 could indicate that ten species are present throughout the country or that twenty species are present in half the area of the country. There are limitations to this index (see Discussion), but by integrating both range area and species richness information, it is more representative of the overall effort a country must give to restore megafauna communities than species richness only. We present results of analysis for MI in the main text of this manuscript and for species richness in Supplementary information 3.

### Socio-economic indicators

We collected socio-economic indicators of countries’ governance, conservation funding and level of human development. Governance consists of the traditions and institutions by which authority in a country is exercised. The Worldwide Governance Indicators (WGI) project reports aggregate and individual governance indicators for over 200 countries and territories over the period 1996–2019, for six dimensions of governance: 1) Voice and Accountability, 2) Political Stability and Absence of Violence, 3) Government Effectiveness, 4) Regulatory Quality, 5) Rule of Law and 6) Control of Corruption (Kaufmann et al. 2010). Together, these indices reflect the capacity of governments to effectively formulate and implement sound policies. We accessed WGI data at info.worldbank.org/governance/wgi/ (accessed 8 October 2020) and extracted values of the 6 indicators for the year 2019. These indices are highly correlated, and we reduced the dataset to one dimension to increase interpretability, using the first component of a principal component analysis (Wold et al. 1987) (Fig S1.1).

Waldron et al. (2013) assembled a large dataset on the financial contributions of countries to conservation that includes both domestic (within-country) spending and donations made by other countries (available at https://datadryad.org/stash/dataset/doi:10.5061/dryad.p69t1, accessed 1 June 2020). The dataset provides mean biodiversity conservation investment for the period 2001-2008 in US 2005 dollars. This represents the latest version of the database that includes both the donor compilation and in countries’ domestic spending (Waldron, pers.comm). We assume that the relative distribution of conservation spending across countries has remained comparable over that period even if absolute levels of conservation spending may have changed since that date. Price level ratio is the ratio of a purchasing power parity (PPP) conversion factor to an exchange rate. It provides a measure of the differences in price levels between countries by indicating the number of units of the common currency needed to buy the same volume of the aggregation level in each country. To account for the difference in overall prices between different countries, we divided dollars spent on conservation in each country by the national price level ratio of the country for the year 2005 (https://data.worldbank.org/indicator/PA.NUS.PPPC.RF, accessed 8 October 2020), following Waldron et al. (2017), to obtain conservation spending values in international dollars. Finally, to make it comparable across countries, we divided conservation spending values by the size of the country. We considered conservation spending per km^2^, hereafter referred to as “conservation funding”, to be representative of the financial ability of countries to apply conservation measures throughout their borders. We excluded 24 countries from the analysis, for which potentially major flows to conservation are unquantified, or where data quality is poor (Waldron et al. 2013, Table S3).

The Human Development Index (HDI) is a composite statistic that is commonly used to indicate human development based on data on education, life expectancy and per capita income. We accessed HDI data for the year 2018 at hdr.undp.org/en/content/human-development-index-hdi (accessed 16 October 2020). Finally, the Organisation for Economic Co-operation and Development (OECD) is an intergovernmental economic organisation, where country members (http://www.oecd.org/about/members-and-partners/) are generally regarded as developed countries with high-income economies and a very high HDI.

### Analyses

We omitted countries that never had megafauna species from the analyses. We tested for a significant difference between MI for OECD vs non-OECD countries for each scenario using Welch’s t-test for samples of unequal variance. To explore the congruence between MI and the financial ability of countries to support conservation, we mapped the combined patterns of MI and socio-economic indices using bivariate choropleth maps. These maps show the relative ranking of countries along two axes, revealing how the indicators combine with each other spatially, and allowing for the identification of spatial clustering. To build these maps, we classified the data in quartiles and assigned a sequential color scheme to the resulting bins. To compare countries’ funding capacity relatively to the rest of the world, we calculated for each country the ratio between total conservation spending (in log) and MI for each scenario. Countries that are overfunded relative to the amount of restoration effort they have to give will have higher values of this ratio, which gives an indication of which countries have a higher capacity to fund megafauna restoration. We compare this ratio to the median ratio for all countries in the world. We also compare the median per continent to the global median.

All analyses were performed in the R programming language v.4.0.2 (R Core Team 2020).

## Results

We consider a total of 147 extant and 151 recently extinct megafauna species. Maps of socio-economic indicators are presented in Supplementary Information 1. Species richness per country ranks from 0 to 38 in the current scenario to 1 to 62 in the Pleistocene scenario (SI, Fig S3.1). Figure 1 shows the global distribution of restoration burden, quantified by MI. This burden is mostly bore by sub-Saharan African countries and south-east Asian countries in the current and historical scenario and increasingly shared with Europe in the Holocene scenario and the Americas in the Pleistocene scenario. This pattern is even more apparent when looking at species richness (SI, Fig S3.1). In a scenario where we restore and protect species within their current range, 19 out of 20 countries with the largest restoration burden are in Africa, with eSwatini, Zambia, Tanzania, Zimbabwe and Kenya having the highest MI scores (Table 1). African countries still bear the highest restoration burden in the historical and Holocene scenario, but an increase in MI is observed in southeast Asia and Europe (Fig 1). If a Pleistocene baseline is used as a benchmark for restoration efforts, new opportunities arise, including in the Global North, with 7 South American countries (Paraguay, Brazil, Suriname, French Guyana, Uruguay, Guyana, Bolivia), 3 European countries (Belarus, Poland, Lithuania) and the United States being part of the 20 countries with the highest MI (Table 2).

**Figure 1.**
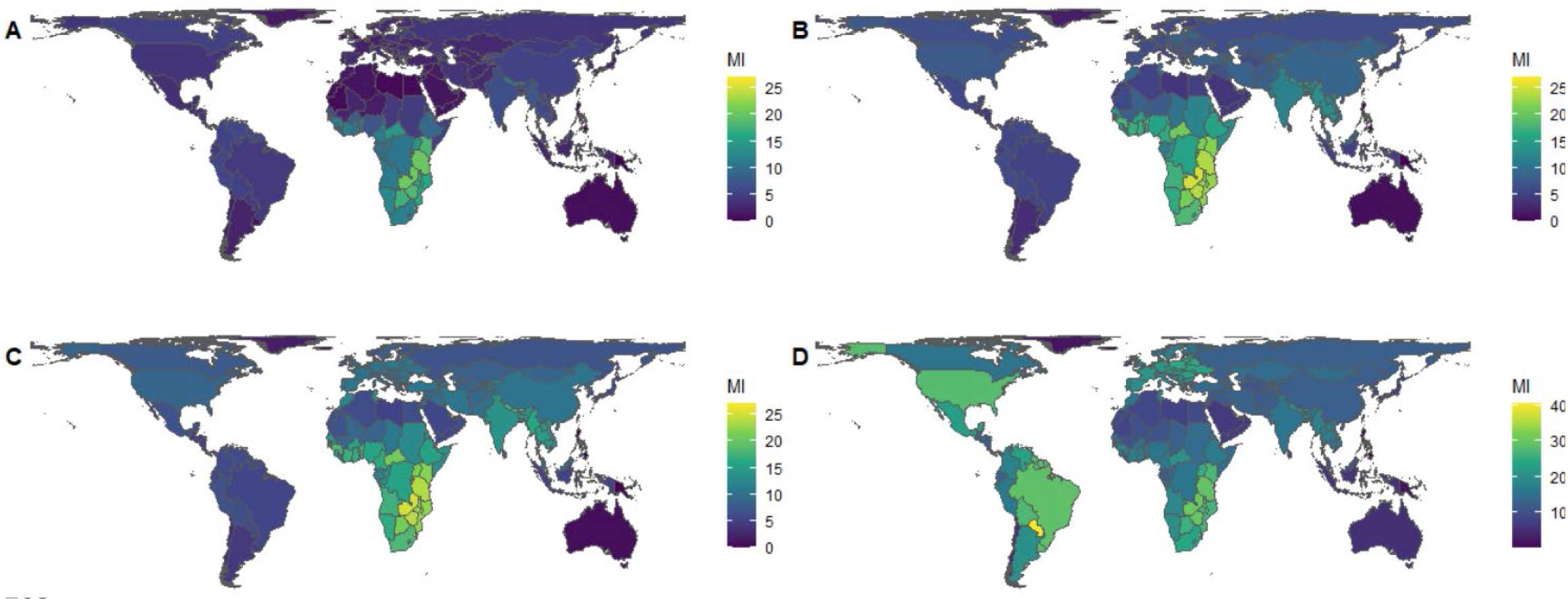
Global distribution of megafauna index (MI) per country for the Current (A), Historical (B), Holocene (C) and Pleistocene (D) scenarios. The scale in panels A-C is the same, to allow for direct comparison.

**Table 1.**
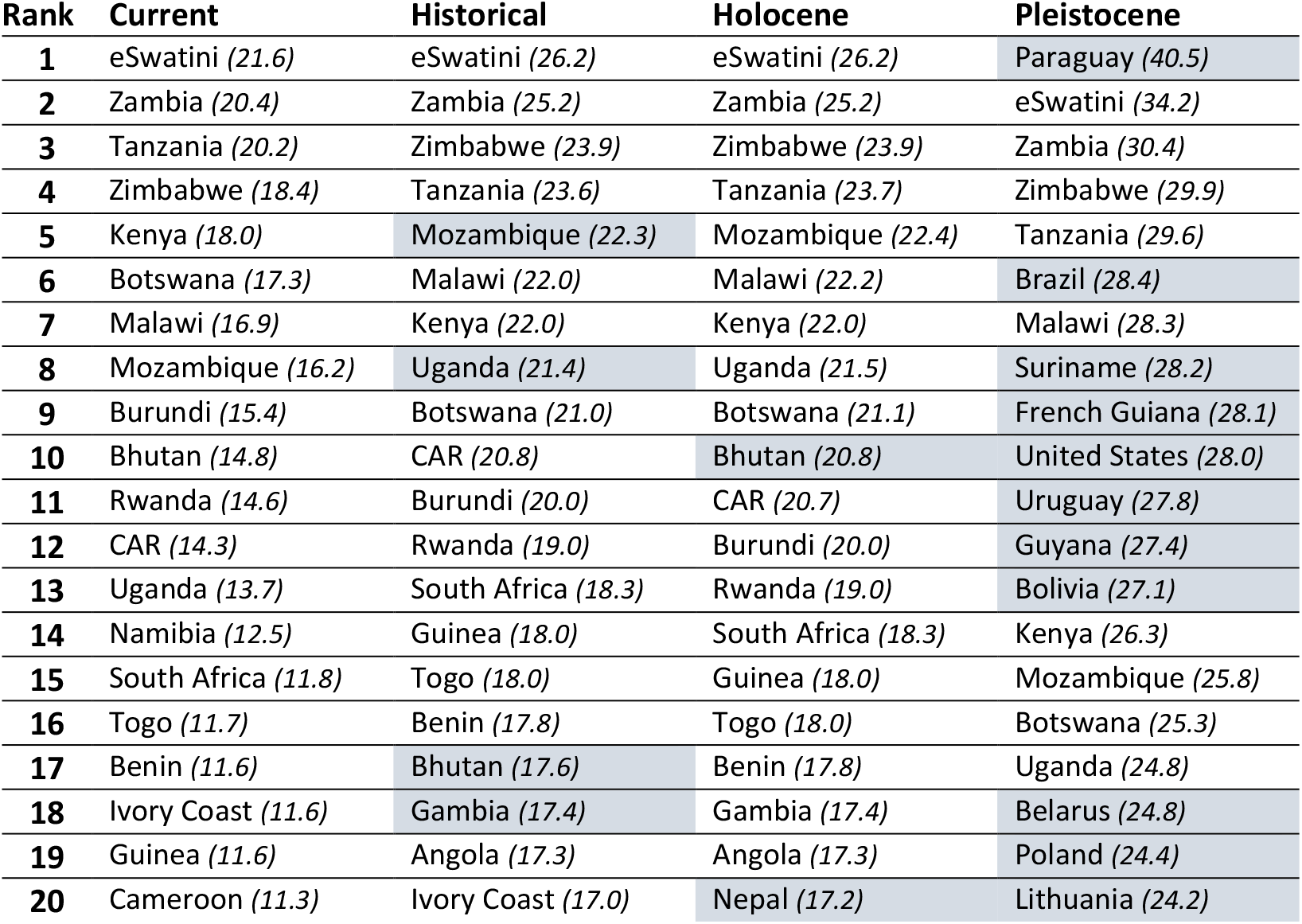
Top 20 countries with the highest megafauna restoration burden for each of the four restoration scenarios. The value of megafauna index is provided in parenthesis. Shaded boxes indicate countries that went up 3 ranks or more compared to the previous scenario. (CAR: Central African Republic)

| Rank | Current | Historical | Holocene | Pleistocene |
| --- | --- | --- | --- | --- |
| 1 | eSwatini (21.6) | eSwatini (26.2) | eSwatini (26.2) | Paraguay (40.5) |
| 2 | Zambia (20.4) | Zambia (25.2) | Zambia (25.2) | eSwatini (34.2) |
| 3 | Tanzania (20.2) | Zimbabwe (23.9) | Zimbabwe (23.9) | Zambia (30.4) |
| 4 | Zimbabwe (18.4) | Tanzania (23.6) | Tanzania (23.7) | Zimbabwe (29.9) |
| 5 | Kenya (18.0) | Mozambique (22.3) | Mozambique (22.4) | Tanzania (29.6) |
| 6 | Botswana (17.3) | Malawi (22.0) | Malawi (22.2) | Brazil (28.4) |
| 7 | Malawi (16.9) | Kenya (22.0) | Kenya (22.0) | Malawi (28.3) |
| 8 | Mozambique (16.2) | Uganda (21.4) | Uganda (21.5) | Suriname (28.2) |
| 9 | Burundi (15.4) | Botswana (21.0) | Botswana (21.1) | French Guiana (28.1) |
| 10 | Bhutan (14.8) | CAR (20.8) | Bhutan (20.8) | United States (28.0) |
| 11 | Rwanda (14.6) | Burundi (20.0) | CAR (20.7) | Uruguay (27.8) |
| 12 | CAR (14.3) | Rwanda (19.0) | Burundi (20.0) | Guyana (27.4) |
| 13 | Uganda (13.7) | South Africa (18.3) | Rwanda (19.0) | Bolivia (27.1) |
| 14 | Namibia (12.5) | Guinea (18.0) | South Africa (18.3) | Kenya (26.3) |
| 15 | South Africa (11.8) | Togo (18.0) | Guinea (18.0) | Mozambique (25.8) |
| 16 | Togo (11.7) | Benin (17.8) | Togo (18.0) | Botswana (25.3) |
| 17 | Benin (11.6) | Bhutan (17.6) | Benin (17.8) | Uganda (24.8) |
| 18 | Ivory Coast (11.6) | Gambia (17.4) | Gambia (17.4) | Belarus (24.8) |
| 19 | Guinea (11.6) | Angola (17.3) | Angola (17.3) | Poland (24.4) |
| 20 | Cameroon (11.3) | Ivory Coast (17.0) | Nepal (17.2) | Lithuania (24.2) |

On average, non-OECD countries have a significantly higher MI than OECD countries for the current, historical and Holocene scenario (Fig 2) (Current: t(171) = 6.43, p<0.001; Historical: t(158) = 5.61, p<0.001; Holocene: t(115) = 3.29, p<0.01). In the Pleistocene scenario, non-OECD countries have a lower restoration burden on average, but the difference is not significant (t(62) = −1.36, p=0.18). We found similar results for species richness (Fig S3.2).

**Figure 2.**
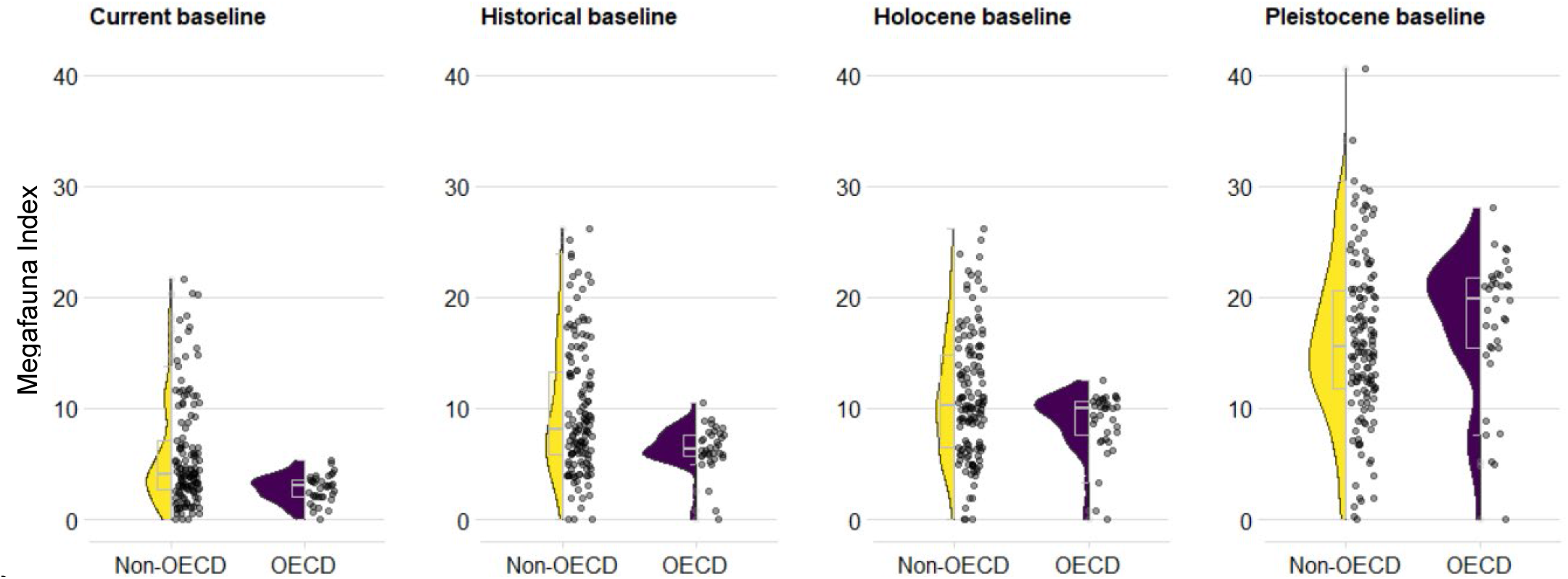
Megafauna index for the four baseline scenarios for OECD (n=36) and non-OECD countries (n=137). Each point represents a country, the violin plot represents the distribution of the data and the grey box plot its summary statistics (median, first and third quartiles). The difference between OECD and non-OECD countries is significant in the three first panels (t-test, p < 0.01) and non-significant in the last panel (Pleistocene, t(62) = −1.36, p=0.18).

Crossing MI with different socio-economic indicators reveals countries that bear a high burden for restoration despite ranking low in terms of governance, conservation funding and HDI (Fig 3, countries in yellow) and *vice versa* (in blue). In the current scenario, most countries fall into one of these two categories, especially for the governance index and HDI (Fig 3, left and right panel). These two indexes are highly correlated (Pearson’s r = 0.8, Fig S1.4) and as a result the spatial pattern is similar. There are a few instances in the current scenario where both MI and the level of conservation funding are high (in purple) e.g., in South Africa, Kenya, India, Thailand and Colombia, but it is important to note that HDI and governance index are low in these countries. In subsequent scenarios, some areas with high values for the three socio-economic indicators also have high values of restoration burden (in purple) e.g., the United States and in western Europe

**Figure 3.**
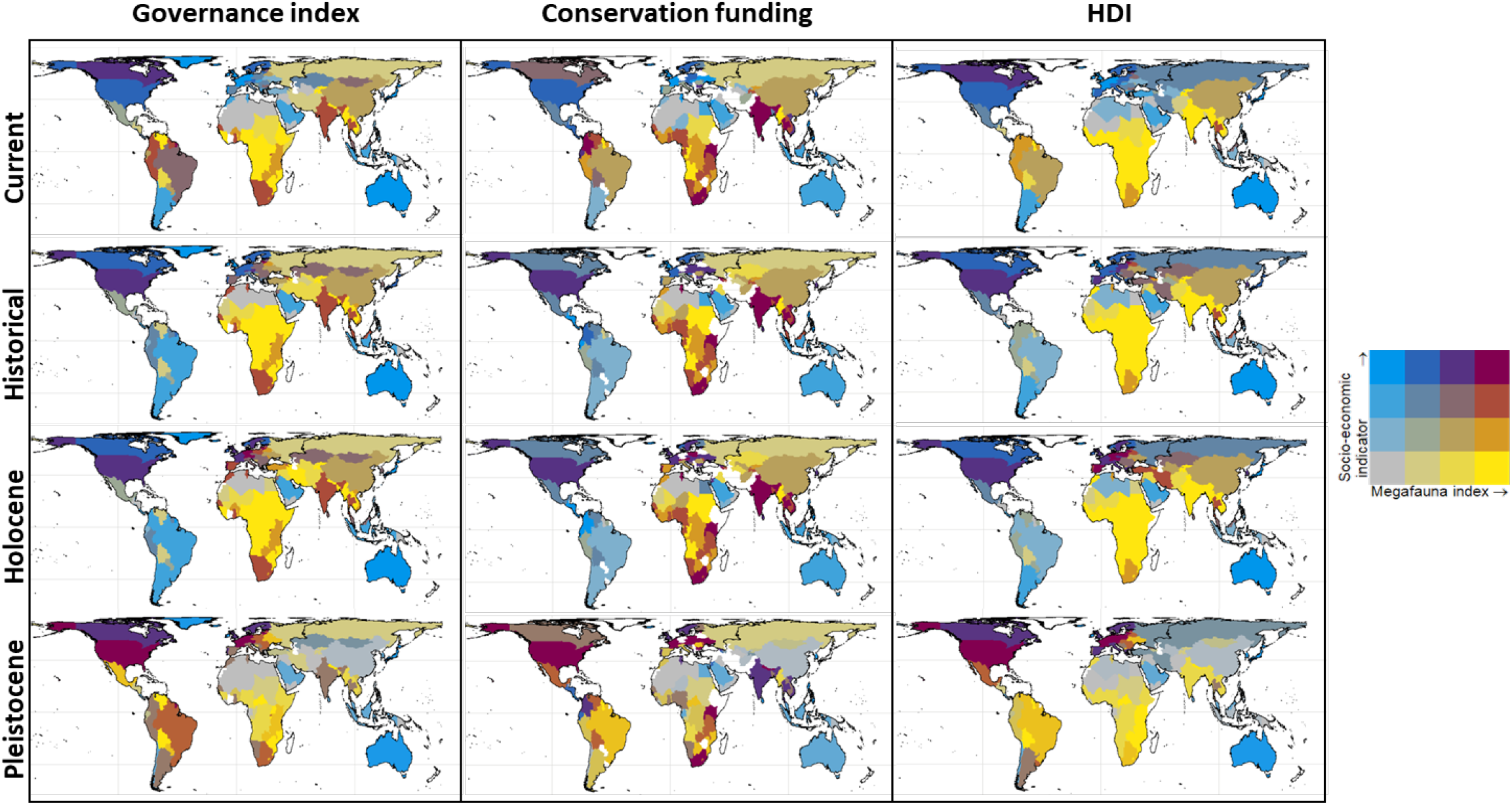
Bivariate choropleth maps of Megafauna Index (MI) versus different socio-economic indicators: Governance index, conservation funding and Human Development Index (HDI), for the four baseline scenarios. See the methods section for a description of the different indices. Values of the indices are divided in quartiles. Yellow colour indicates a high MI but low values for the given socio-economic indicator, and vice versa for blue colours. Purple indicates that the level of MI is commensurate with the value of socio-economic index in the country, relatively to the rest of the world. Countries in white are those that never had megafauna species. In addition, 24 countries were excluded from the analysis of conservation funding (middle panel) due to poor data quality (see Methods).

Additional analyses show a negative correlation between MI and HDI for the current (Pearson’s r = −0.57, p < 0.001), historical (r = −0.62, p < 0.001) and Holocene (r = −0.52, p < 0.001) scenarios, which is not significant in the Pleistocene scenario (r = −0.04, p > 0.05) (Supplementary Material, Fig S2.1, Table S2.1). A negative correlation is also observed for governance index for the current (Pearson’s r = −0.31, p < 0.001), historical (r = −0.35, p < 0.001) and Holocene (r = −0.26, p < 0.001) scenarios, and not in the Pleistocene scenario (r = −0.07, p > 0.05) (Supplementary Material, Fig S2.2, Table S2.2). We did not find any significant correlation between MI and conservation funding for the four scenarios (Supplementary Material, Fig S2.3, Table S2.3).

Finally, Fig. 4 shows the ratio between conservation funding and MI, indicating how countries are under- or over-funded relative to the rest of the world, given the value of MI. Countries on the right of the vertical bar (which represents the global median value of the ratio) are relatively over-funded relative to the rest of the world (and *vice versa* for countries on the left). These results suggests that African countries are generally underfunded for megafauna restoration compared to the rest of the world for all scenarios. All North American countries, Australia and most European countries are consistently overfunded in all scenarios, though the distance to the global median is smaller in the Pleistocene scenario. In that scenario, South American countries are on average underfunded for megafauna restoration.

**Figure 4.**
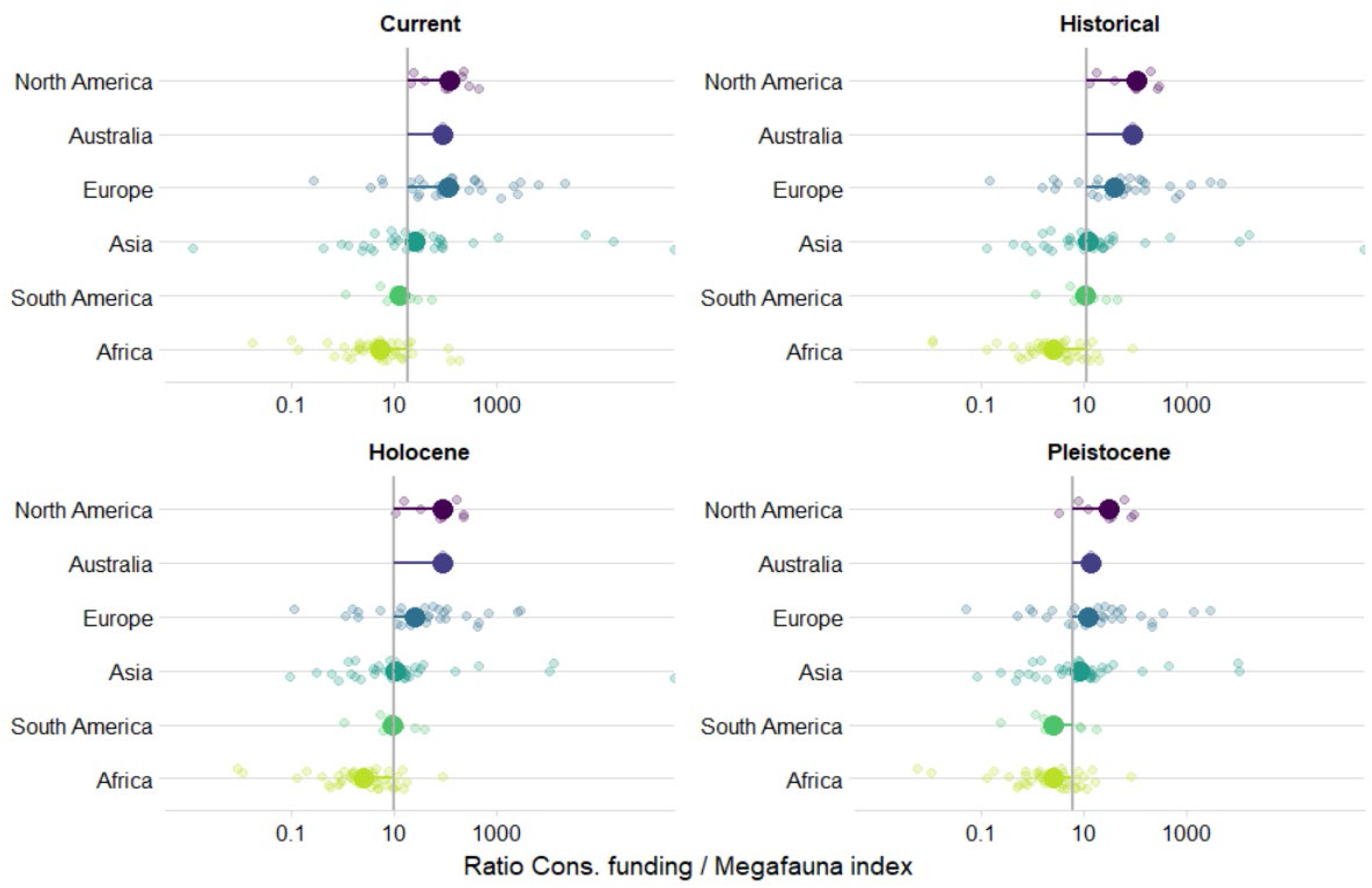
Ratio between conservation spending and megafauna index (MI) per country, grouped by continent, and for the four baseline scenarios. A higher ratio represents a high conservation capacity in terms of funding, relative to the value of MI. Each small point represents a country and large points are the median value of the ratio for the whole continent. In each panel, the vertical bar corresponds to the global median value of the ratio for that scenario. Countries on the left of that bar are relatively under-funded compared to the rest of the world, while countries on the right of the bar are relatively over-funded. Note that the data has been log-transformed and the differences observed on the graph are thus exponential.

## Discussion

In his plead for a full reckoning of the historical biogeography of species in conservation, Sanderson (2019) argues that *“picking a particular date [to define the indigenous range of a species] means arbitrarily accepting some events as incontrovertible while requiring others to be addressed through conservation action”,* and that using recent or historical baselines *“stamps conservation with a Eurocentric perspective that is unwarranted for a global enterprise”.* This post-Columbian bias has been highlighted and discussed in the context of rewilding before (Donlan et al. 2006, Keulartz 2016, Svenning and Faurby 2017, Jordan et al. 2020), but we for the first time quantify these global geographic patterns for megafauna restoration, linking them to contemporary measures of human development, funding and governance. Our analysis highlights the uneven distribution of the burden of megafauna restoration across the world, and how this geographic pattern changes with the benchmark used to guide restoration. We identify areas where high levels of megafauna restoration index overlap with lower financial and societal capacity to support megafauna conservation and restoration efforts.

Our results show that using a recent or historical (1500 AD) temporal baseline as a benchmark for restoration puts higher responsibility for restoring megafauna communities on countries in the Global South, which also rank lower in conservation funding, governance and human development. African countries in particular bear a high burden for megafauna restoration, being the continent where less megafauna species went extinct in the Late Pleistocene (Smith et al. 2018), and which is home to some of the greatest diversity of extant megafauna today (Ripple et al. 2016). The burden for megafauna restoration in countries in the Global North increases when a Holocene (for Eurasia) or Pleistocene (for North America) baseline is used as a benchmark for restoration, thus reducing the current observed negative correlation between restoration burden and socio-economic indicators. High levels of funding ability, governance and human development would make North America and Europe priority areas for rewilding if a Pleistocene or Holocene benchmark is used. Overall, these results highlight how the identification of global priority areas for ecosystem restoration using recent benchmarks may put an unfair burden on areas where impacts on biodiversity were the most recent and underestimate the restoration potential in other.

### Limitations

Aiming for a comprehensive characterization of the trajectories of governments, ecosystems, and the Earth system in relation to trophic rewilding is beyond the scope of this paper. Instead, this study uses available data and simple tools to explore scenarios of restoration and discuss emerging patterns. This creates certain limitations in the interpretation and application of these results.

First, the analysis assumes that species are being reintroduced throughout their range, which does not address the potential ecological, practical, social, and political barriers to megafauna restoration (Lorimer et al. 2015). Rather than providing realistic spatial prioritization for trophic rewilding, this analysis aims to stimulate a debate based on the representation of alternative environmental futures. Successful restoration of megafauna will depend on tolerant socio-political landscapes and favourable ecological conditions (Treves and Karanth 2003), which are better explored with conservation planning tools and taking into account socio-ecological frameworks (Wilson et al. 2009, Ceauşu et al. 2019, Thierry and Rogers 2020), while being mindful of ethical considerations for both the animals and humans involved (Thulin and Röcklinsberg 2020). Climate change will also provide its own sets of challenges for nature conservation politics (Titley et al. 2021) and projecting potential range shifts under future climate is a priority for future research.

Second, MI does not differentiate between species, resulting in a similar burden for species that may have very different impacts on society and requirements for their conservation. This can potentially affect results’ interpretation if distribution patterns of the most challenging species to conserve, e.g. largest body-sized carnivores and herbivores (Rabinowitz 2005, Thirgood et al. 2005, Gulati et al. 2021), are strongly biased towards certain countries. This may be true for the current and historical scenarios, where high body-mass and large carnivore species are mostly found in Africa and southeast Asia. These countries may thus bear an even higher burden for megafauna restoration than presented in this study, which makes our interpretations conservative. Plus, the index we use in this study includes information on within-country range size, which correlates to body mass (Kelt et al. 2001), thus indirectly including some aspects of species’ size in the calculation of MI and potentially reducing this bias.

Finally, we acknowledge the subjective value system with which we approach the issue of megafauna restoration. While the term “burden” has been used before to refer to megafauna conservation (see e.g. Lindsey et al. 2017, Flannery and Boitani 2018 p. 175, Jordan et al. 2020), it introduces a biased perception of megafauna which does not recognize the multiplicity of – potentially positive – experiences that different stakeholders may have with these species (Clayton 2019). In what we call a burden, some might see an opportunity, e.g. for improved livelihoods through employment in nature-based tourism and the provision of associated goods and services (Navarro and Pereira 2015, Willemen et al. 2015), economic advantages for land management through wilder landscapes requiring less–or more adaptive–management than conventional alternatives (Navarro and Pereira 2015), or for more functional ecosystems (Perino et al. 2019, Schowanek et al. 2021). Some may also recognize the intrinsic value of megafauna, independent of human uses and valuation (Lockwood 1999). This paper does not pretend to delve into the complexities of how megafauna are perceived in different contexts and by a range of actors. By using the term “burden”, we attract the reader’s attention towards the potential costs of managing and coexisting with these species that are often disproportionately borne by institutions and local rural communities in the Global South (Jordan et al. 2020).

### Recommendations

With these limitations in mind, we hope to contribute to the debate around the choice of benchmarks for restoration, by highlighting the need for a mindful evaluation of restoration potential patterns and the distribution of financial and socio-political resources across the world. To achieve a more equitable distribution of the burden of restoration, we suggest 1) increased support from the Global North to build capacity in areas where the ratio of cost to benefit for megafauna restoration is high and 2) recognition of the shared responsibility that both the North and the South have in preserving and restoring large mammal communities for the future.

As suggested in the literature and highlighted in this study, the focus on African and south-eastern Asian countries for megafauna conservation has mostly been reinforced by the tendency of conservation biologists to consider recent or pre-European baselines as benchmarks for restoration, overlooking the potential for restoration in areas with a long history of environmental degradation (Donlan et al. 2006, Jordan et al. 2020). Financial mechanisms are already in place to ensure a transfer of foreign aid to those countries who are underfunded for conservation (Miller et al. 2013, Waldron et al. 2017). However, they fall short of meeting conservation needs for megafauna today (Waldron et al. 2013, Lindsey et al. 2017) and this trend is likely to accentuate if countries have to move beyond conservation and also actively restore megafauna species within their historical range. Meeting the costs of megafauna restoration globally will require substantial transfer of resources from the Global North to the Global South (Balmford and Whitten 2003), and appropriate support from local governments and the international community to build and maintain appropriate levels of conservation capacity (O’Connell et al. 2019).

Countries from the North benefit from the global ecosystem services, existence values and direct use values associated with conservation in other countries without incurring the costs (Balmford et al. 2003). There is thus a certain hypocrisy with richer countries expecting low-income countries to demonstrate greater tolerance to coexistence with megafauna at the same time as they often struggle to apply the same principles within their borders (Jordan et al. 2020). Lions, rhinos and elephants were part of European landscapes until at least the Late Pleistocene and in North America, just 13,000 years ago, fifty additional megafauna species were roaming in the landscape, including mammoths, mastodons, horses, ground sloths, camels, dire wolves, lions and sabertooth cats (Stock 1930). The Americas and Australia have suffered severe megafauna extinctions and have few remaining candidate species for restoration. However, ambitious restoration approaches could participate in recovering Pleistocene-levels of diversity and aim towards ambitious restoration targets (Seddon et al. 2014). A subset of these extinct species have extant functional analogues – closely related taxa with similar functions in the ecosystem – or naturalized nonnative populations that could be used as surrogates in restoration programs (Donlan et al. 2006, Lundgren et al. 2020). This aspect remains controversial (Oliveira-Santos and Fernandez 2010), as illustrated by strong negative public reactions to the idea of North American Pleistocene rewilding (Donlan and Greene 2010), and requires careful risk-assessment. North Africa, Europe and Asia have a high potential for rewilding as functional assemblages could be partially restored using only native species (Schowanek et al. 2021). There, the restoration of extant species into their original geographic distributions (our Holocene baseline) is a more attainable goal and could be a first step towards reversing long-term defaunation in the Global North. It does not come without challenges, as highlighted by public resistance to the comeback of large carnivores (Knight 2016, Behr et al. 2017, Kuijper et al. 2019) or even beavers (Coz and Young 2020) in Europe. But extant megafauna are already making a comeback, including in Europe (Deinet et al. 2013, Chapron et al. 2014), highlighting the potential for continental-scale conservation of large mammals in human-dominated landscapes (Linnell et al. 2020). With this long-term perspective, restoring megafauna is not less relevant in the Global North and also has the potential to produce large benefits for conservation and restoration (Svenning 2020, Schowanek et al. 2021).

### Baselines in restoration

The selected reference state to define species’ indigenous range will always influence the magnitude of damage measured, which becomes vitally important when setting quantitative targets for restoration and measuring success towards recovery. Ultimately, the choice of a given reference state is always situated within larger socio-cultural contexts and arguments over baselines may become arbitrary (Keulartz 2016). The interpretation of reference states should be tailored to specific contexts, spatial and temporal scales (McNellie et al. 2020). For example, while Pleistocene rewilding can be met with criticism (Rubenstein et al. 2006, Oliveira-Santos and Fernandez 2010), this long-term perspective is useful to provide context for future-oriented restoration initiatives, for informing on the potential for biodiversity (in contrast to more degraded Holocene time points) and for informing on the factors that have been key for generating and maintaining biodiversity in the long-term (Svenning 2020). Holocene baselines also provide invaluable insights into species’ ecology, climatic tolerance and habitat preferences that are often not available from recent records only (Monsarrat et al. 2019).

Historical conditions may be an unattainable reference state for contemporary ecosystems (McNellie et al. 2020), which led some conservationists to suggest a future-oriented reference state for rewilding, promoting novel or no-analog ecosystems that may contain new, non-historical combinations of species (Hobbs et al. 2009). While this framework aims to move away from historical perspectives, it still relies on the identification of areas where targeted restoration actions might lead to the return of historically extirpated species, or even to novel colonists (Mason et al. 2021). This approach thus calls for a reckoning of the past to understand where species would likely exist today in the absence of historic human impacts. It also benefits from better understanding of the factors and processes that have generated and maintained species richness in the long-term, which are provided by studies taking a deep-time perspective (Svenning 2020).

A reflexive approach on the value system by which conservation biologists and stakeholders define baselines and consequently classify species as native is key to challenge our perception of what we perceive as possible today and in the future. We illustrate this with the example of megafauna species and rewilding. However, long-term human impacts have affected a wide range of species and restoration of any group of taxa that have undergone range contraction in the past will pose similar issues. We suggest that real transformation will only be achieved if wealthier countries take responsibility for achieving global restoration objectives, both by supporting restoration efforts in the South and in their own backyard.

## Data archiving statement

Species’ current and present-natural ranges from the PHYLACINE database are openly available on github (https://github.com/MegaPast2Future/PHYLACINE_1.2). Datasets of socio-economic indicators used in the analyses are available at https://data.worldbank.org/indicator and http://hdr.undp.org/ and conservation spending data at https://datadryad.org/stash/dataset/doi:10.5061/dryad.p69t1. The data and R code to reproduce the findings of this study are openly available in Figshare https://figshare.com/s/8312ba0f8a9d446f77de.

## Acknowledgements

We thank Anthony Waldron for his advice on the conservation funding dataset. We also thank Robert Buitenwerf and other members of the MegaPast2Future group for constructive feedback on earlier versions of these analyses, and Ashley Pearcy Buitenwerf for feedback on the text.

