## Supplementary Information for "Using recent baselines as benchmarks for megafauna restoration places an unfair burden on the Global South"

### The burden of megafauna restoration varies across the world when different temporal baselines are used as benchmarks for restoration

---

<u>Supplementary Information 1:</u>

Socio-economic indicators

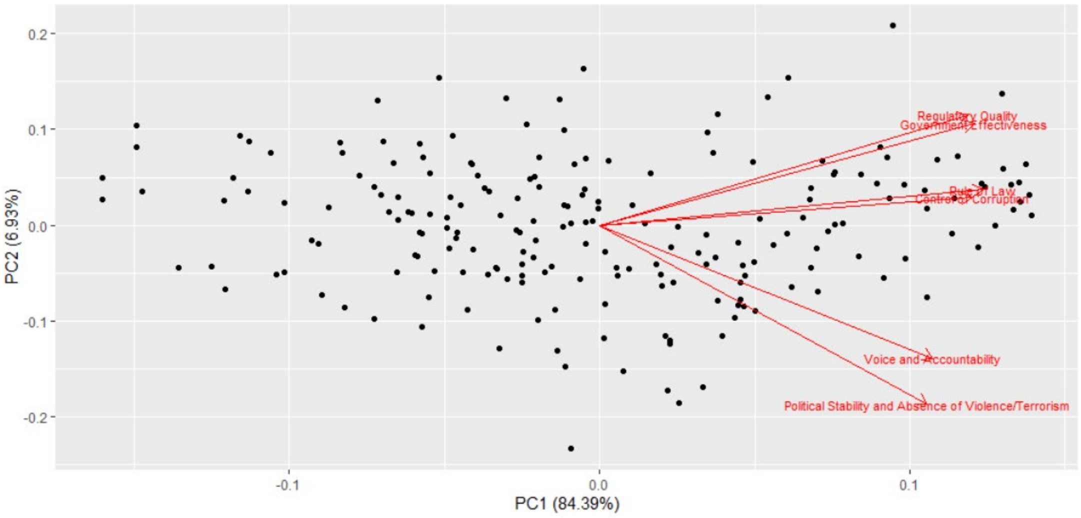


***Figure S1.1. Principal Component Analysis on the 6 dimensions of governance indicators** (Voice and Accountability, Political Stability and Absence of Violence, Government Effectiveness, Regulatory Quality, Rule of Law, Control of Corruption [http://info.worldbank.org/governance/wgi/](http://info.worldbank.org/governance/wgi/)).*

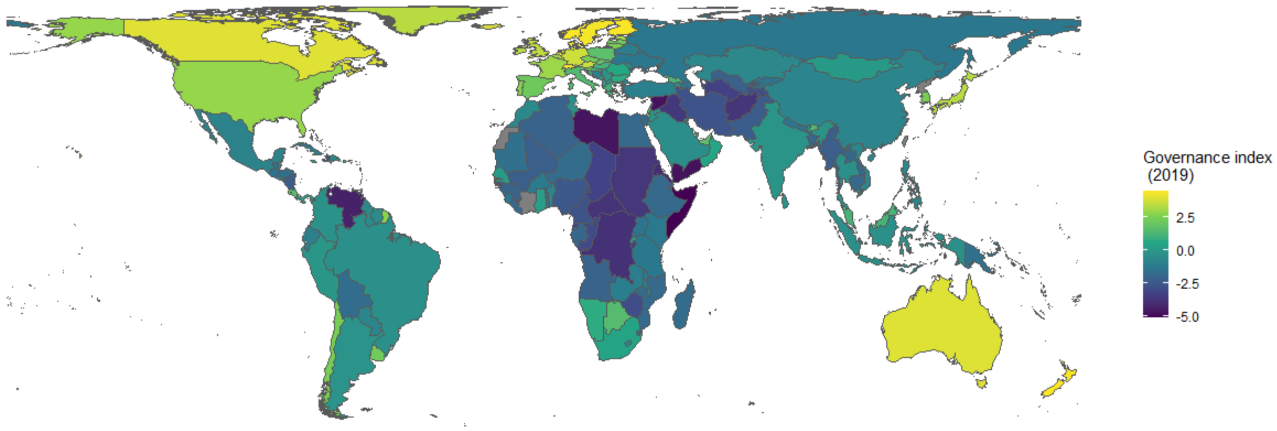


***Fig S1.2. Global distribution of the governance index used in this study** (see Methods in the main text for a description of the index).*

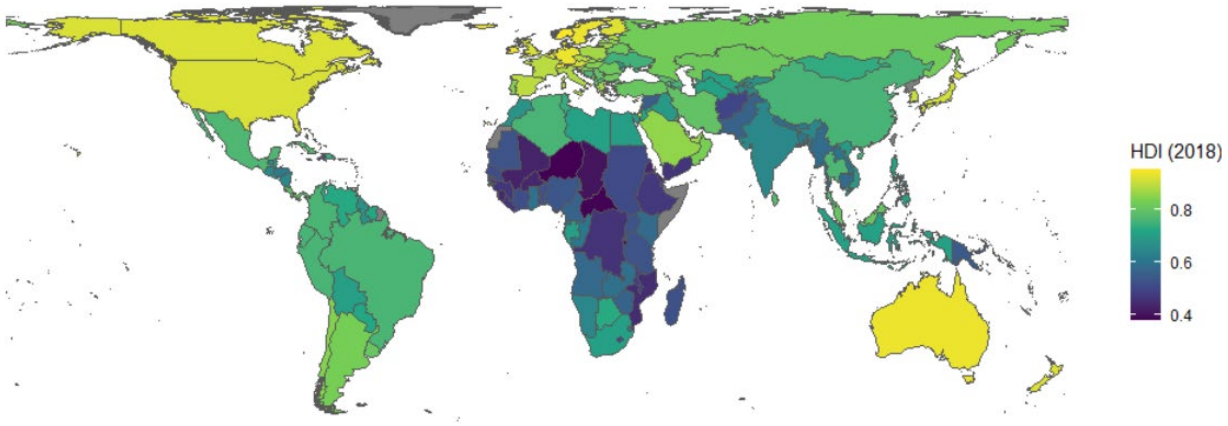


***Figure S1.3. Global distribution of Human Development Index 2018*** *(see Methods in the main text for a description of the index).*

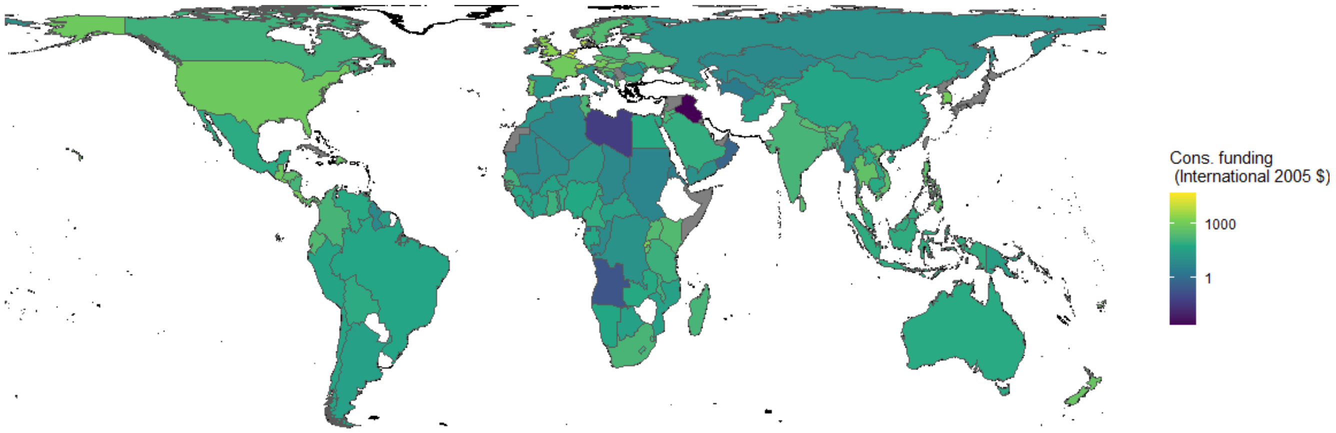


***Fig S1.3. Global distribution of conservation funding index (in International 2005 dollars per km²).***
*Countries in white were excluded from the analysis of conservation funding (middle panel) due to poor data quality (see Methods in the main text for a description of the index).*

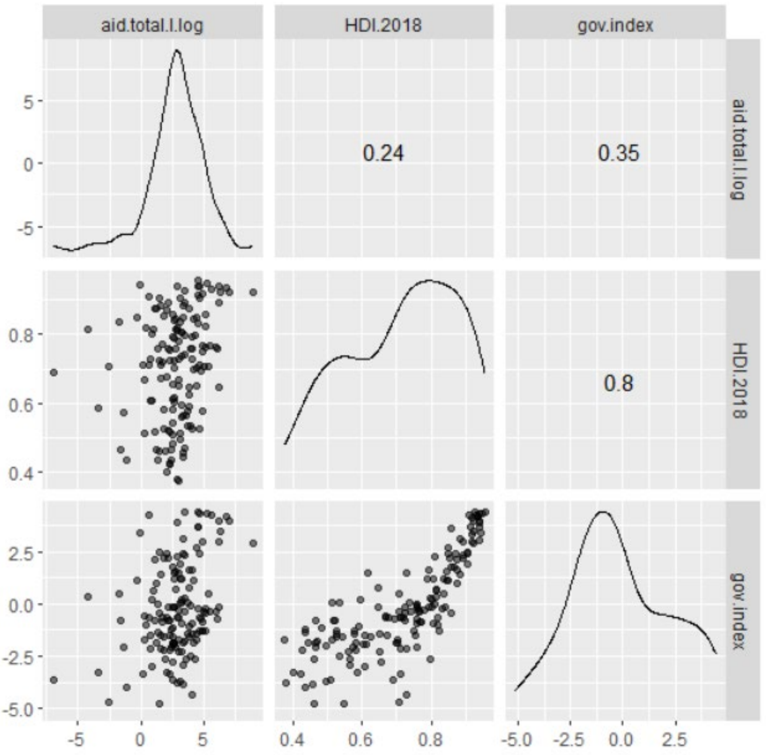


***Figure S1.4. Scatterplot matrix, showing density plots and Pearson's r correlation value, for the three socio-political indexes used in the analysis*** *- Conservation funding (aid.total.I.log), HDI (HDI.2018) and governance (gov.index).*

<u>Supplementary Information 2:</u>

Scatter plots and correlation tests of megafauna index vs socio-economic indicators

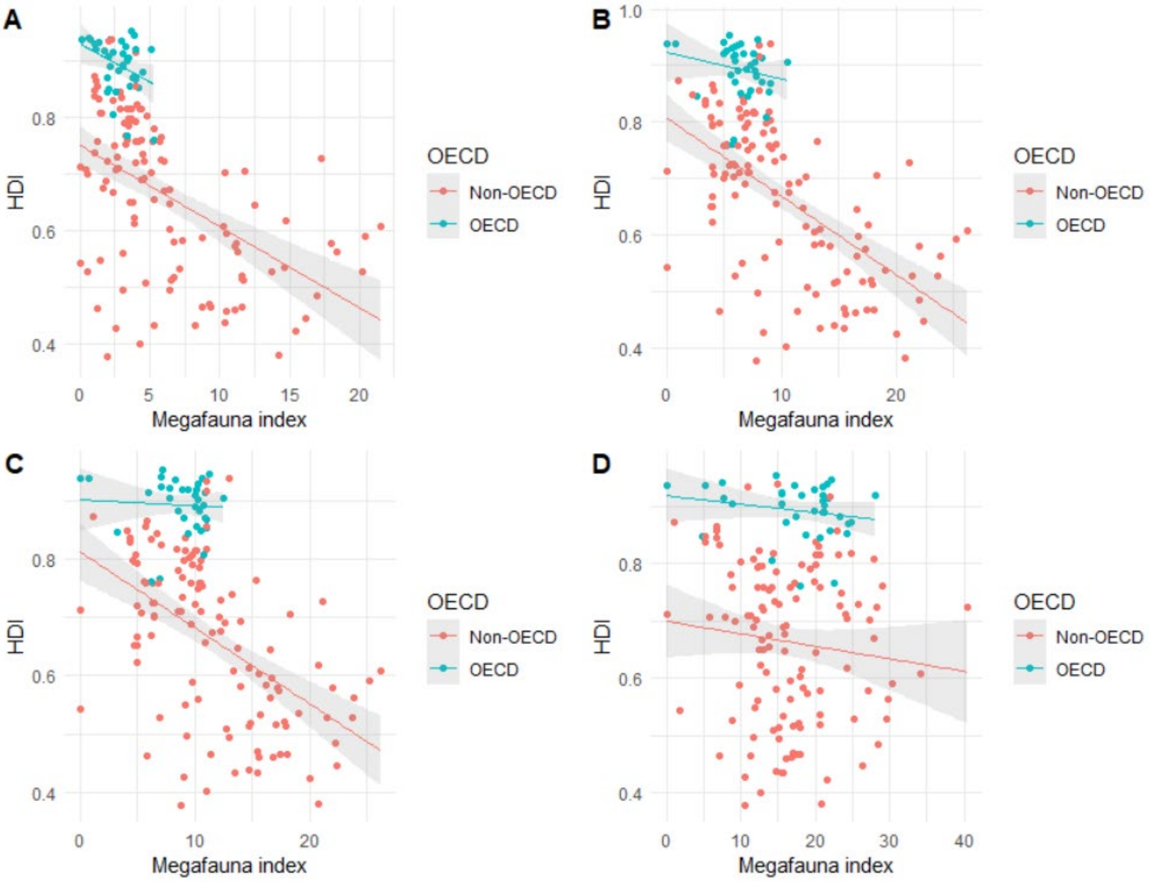


***Fig S2.1. Scatter plots of megafauna index vs the Human Development Index (HDI) for the four restoration scenarios*** *A) Current baseline, B) Historical baseline, C) Holocene Baseline, D) Pleistocene baseline. See main text for a description of this indices. Country members of the OECD are represented in blue and other countries in pink. The lines and shaded areas represent the linear smoother of the data and 95% confidence intervals.*

***Table S2.1.*** *Pearson correlation coefficient and statistical significance levels of megafauna index vs HDI indicators for the four restoration scenarios (*p < 0.05, ***p<0.001).*

| | **Non-OECD** | **OECD** | **Total** |
|---|---|---|---|
| **Current** | -0.51*** | -0.36* | -0.57*** |
| **Historical** | -0.59*** | -0.21 | -0.62*** |
| **Holocene** | -0.52*** | -0.06 | -0.52*** |
| **Pleistocene** | -0.11 | -0.20 | -0.04 |

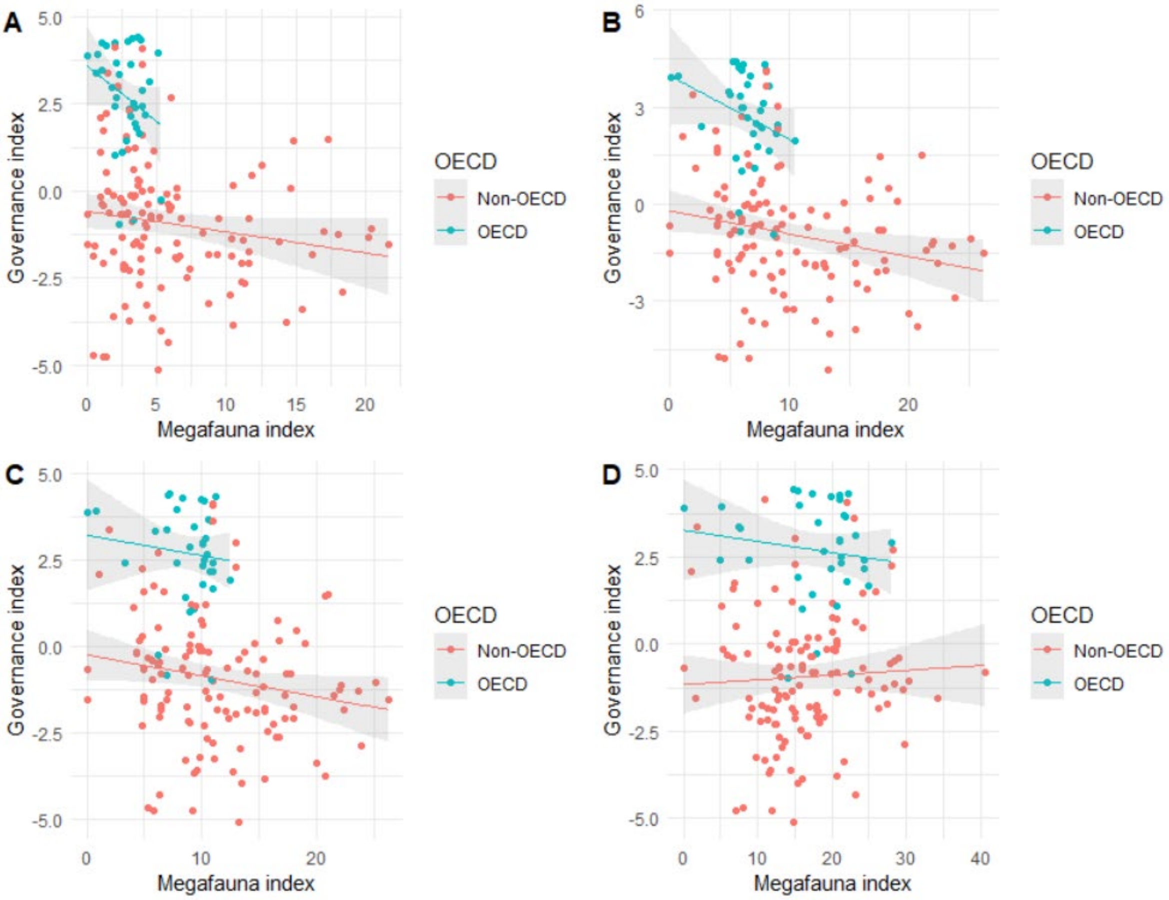


***Fig S2.2. Scatter plots of megafauna index vs governance index for the four restoration scenarios*** *A) Current baseline, B) Historical baseline, C) Holocene Baseline, D) Pleistocene baseline. See main text for a description of this indices. Country members of the OECD are represented in blue and other countries in pink. The lines and shaded areas represent the linear smoother of the data and 95% confidence intervals.*

***Table S2.2.*** *Pearson correlation coefficient and statistical significance levels of megafauna index vs governance index for the four restoration scenarios (*p < 0.05, ***p<0.001).*

| | **Non-OECD** | **OECD** | **Total** |
|---|---|---|---|
| **Current** | -0.16 | -0.28 | -0.31*** |
| **Historical** | -0.22* | -0.29 | -0.35*** |
| **Holocene** | -0.18* | -0.12 | -0.26*** |
| **Pleistocene** | 0.05 | -0.14 | 0.07 |

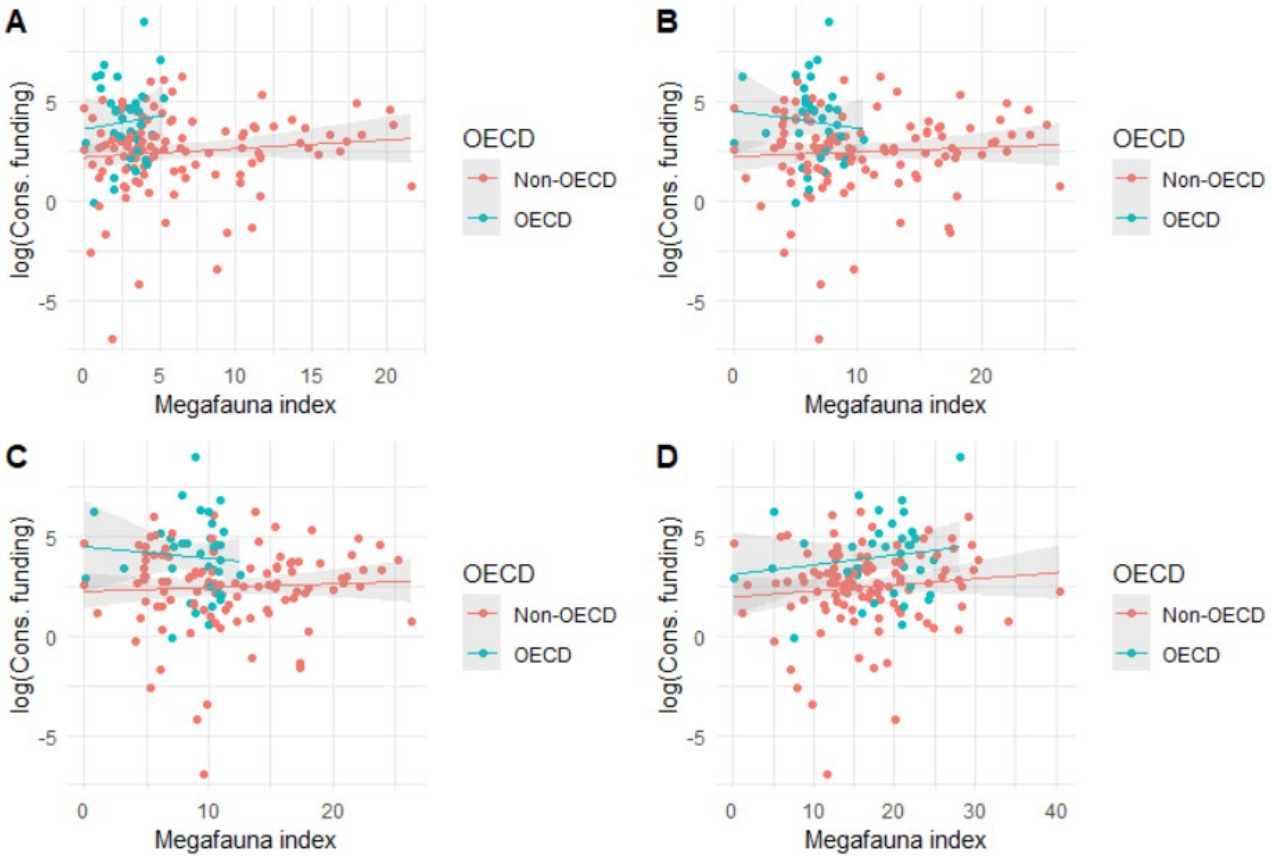


***Fig S2.3. Scatter plots of megafauna index vs log-transformed conservation funding for the four restoration scenarios*** *A) Current baseline, B) Historical baseline, C) Holocene Baseline, D) Pleistocene baseline. See main text for a description of this indices. Country members of the OECD are represented in blue and other countries in pink. The lines and shaded areas represent the linear smoother of the data and 95% confidence intervals.*

***Table S2.3.*** *Pearson correlation coefficient and statistical significance levels of megafauna index vs conservation funding for the four restoration scenarios.*

| | **Non-OECD** | **OECD** | **Total** |
|---|---|---|---|
| **Current** | 0.11 | 0.08 | -0.01 |
| **Historical** | 0.07 | -0.09 | -0.05 |
| **Holocene** | 0.05 | -0.08 | -0.03 |
| **Pleistocene** | 0.11 | 0.15 | 0.14 |

Supplementary Information 3:

Analysis of species richness

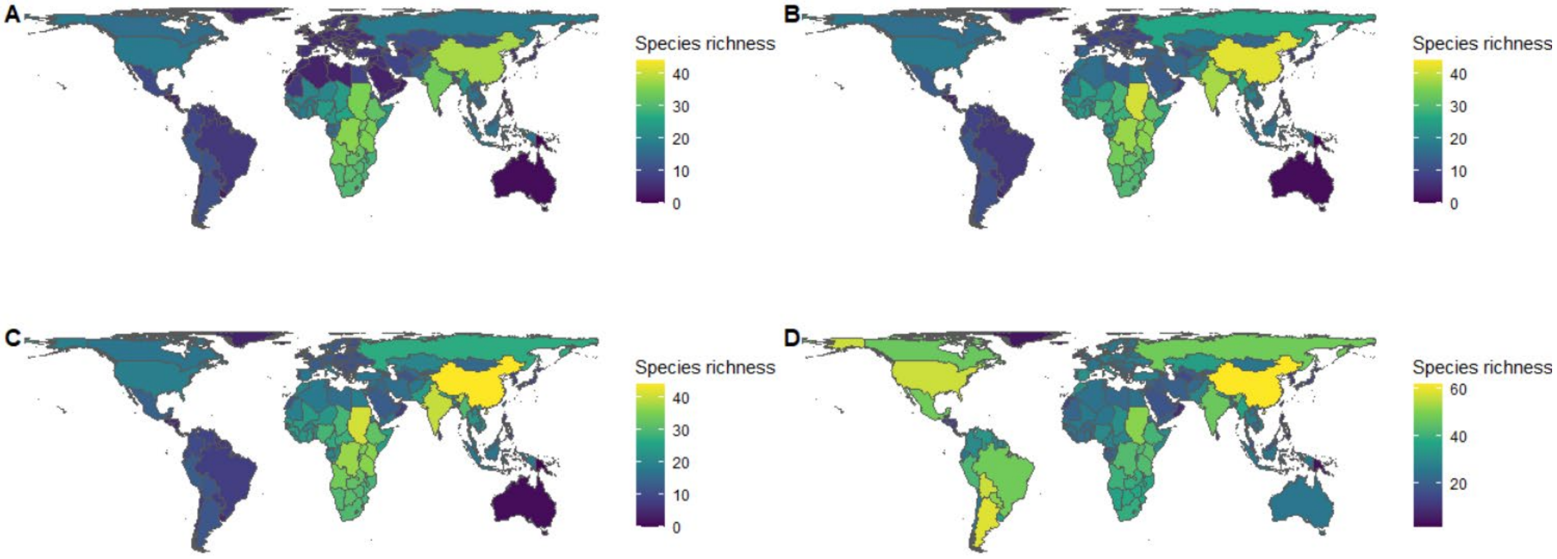


***Figure S3.1.*** *Global distribution of megafauna species richness country for the Current (A), Historical (B), Holocene (C) and Pleistocene (D) scenarios. The scale in panels A-C is the same, to allow for direct comparison.*

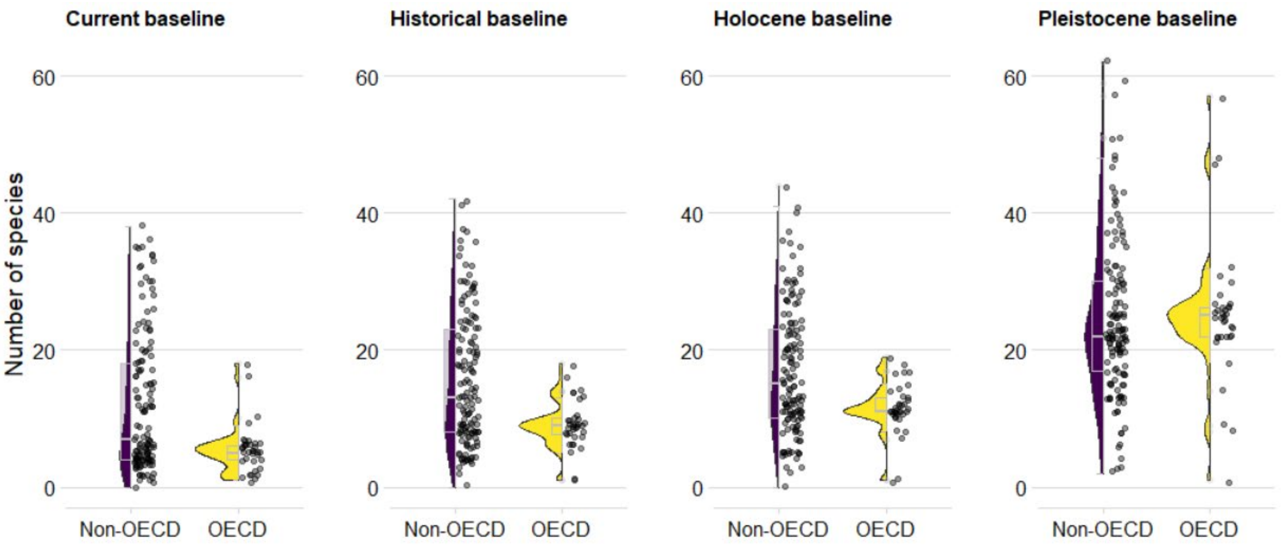


***Figure S3.2. Species richness for the four baseline scenarios for OECD (n=36) and non-OECD countries (n=137).*** *Each point represents a country, the violin plot represents the distribution of the data and the grey box plot its summary statistics (median, first and third quartiles). The difference between OECD and non-OECD countries is significant in the three first panels (t-test, p < 0.001) and non-significant in the last panel (Pleistocene, t(62) = -0.2973, p=0.77).*

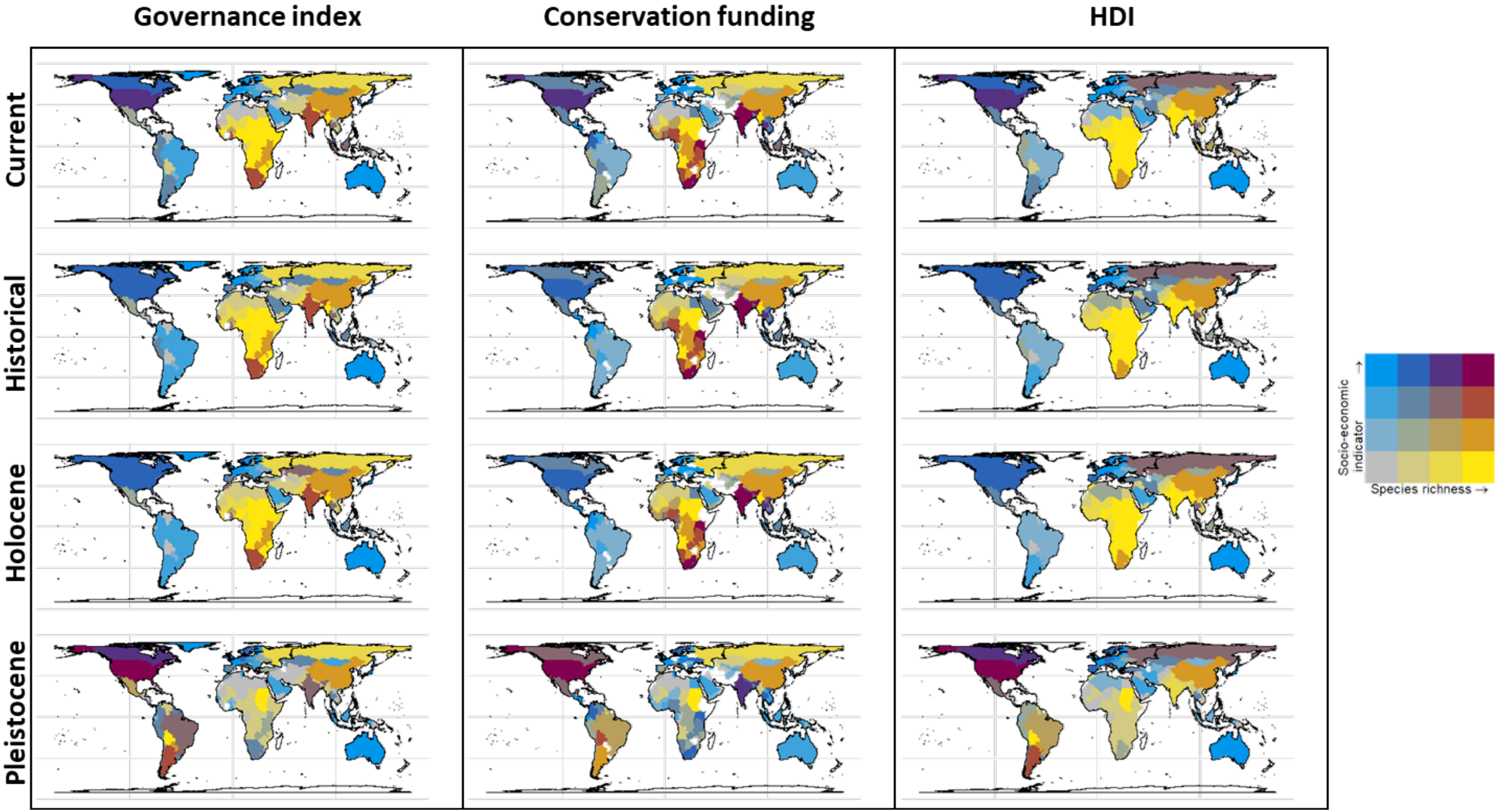


***Figure S3.3.*** *Bivariate choropleth maps of species richness versus different socio-economic indicators: Governance index, conservation funding and Human Development Index (HDI), for the four baseline scenarios. See the methods section in the main text for a description of the different indices. Values of the indices are divided in quartiles. Yellow colour indicates a high number of species but low values for the given socio-economic indicator, and vice versa for blue colours. Purple indicates that the level of MI is commensurate with the value of socio-economic index in the country, relatively to the rest of the world. Countries in white are those that never had megafauna species. In addition, 24 countries were excluded from the analysis of conservation funding (middle panel) due to poor data quality (see Methods in main Text).*

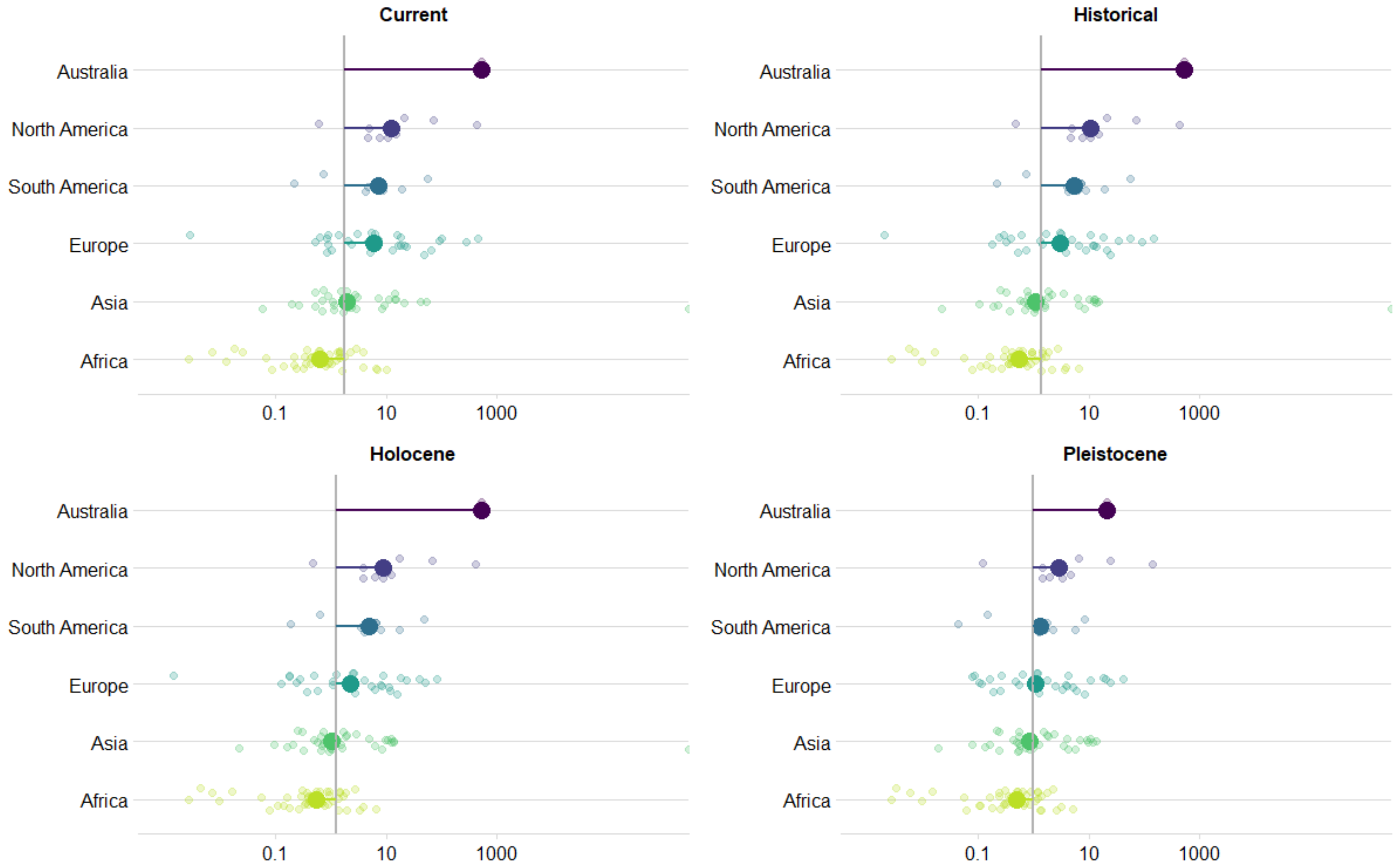


***Figure S3.4.*** *Ratio between conservation spending and species richness per country, grouped by continent, and for the four baseline scenarios. A higher ratio represents a high conservation capacity in terms of funding, relative to the number of species to restore. Each small point represents a country and large points are the median value of the ratio for the whole continent. In each panel, the vertical bar corresponds to the global median value of the ratio for that scenario. Countries on the left of that bar are relatively under-funded compared to the rest of the world, while countries on the right of the bar are relatively over-funded. Note that the data has been log-transformed and the differences observed on the graph are thus exponential.*
